# Whole-genome long-read sequencing downsampling and its effect on variant calling precision and recall

**DOI:** 10.1101/2023.05.04.539448

**Authors:** William T. Harvey, Peter Ebert, Jana Ebler, Peter A. Audano, Katherine M. Munson, Kendra Hoekzema, David Porubsky, Christine R. Beck, Tobias Marschall, Kiran Garimella, Evan E. Eichler

## Abstract

Advances in long-read sequencing (LRS) technology continue to make whole-genome sequencing more complete, affordable, and accurate. LRS provides significant advantages over short-read sequencing approaches, including phased *de novo* genome assembly, access to previously excluded genomic regions, and discovery of more complex structural variants (SVs) associated with disease. Limitations remain with respect to cost, scalability, and platform-dependent read accuracy and the tradeoffs between sequence coverage and sensitivity of variant discovery are important experimental considerations for the application of LRS. We compare the genetic variant calling precision and recall of Oxford Nanopore Technologies (ONT) and PacBio HiFi platforms over a range of sequence coverages. For read-based applications, LRS sensitivity begins to plateau around 12-fold coverage with a majority of variants called with reasonable accuracy (F1 score above 0.5), and both platforms perform well for SV detection. Genome assembly increases variant calling precision and recall of SVs and indels in HiFi datasets with HiFi outperforming ONT in quality as measured by the F1 score of assembly-based variant callsets. While both technologies continue to evolve, our work offers guidance to design cost-effective experimental strategies that do not compromise on discovering novel biology.

## INTRODUCTION

Over the last five years, long-read sequencing (LRS) technologies have transformed the landscape of genetic variant discovery in two fundamental ways. First, they have increased the sensitivity of structural variant (SV) discovery by approximately threefold by providing access to repetitive regions of genomes typically masked or excluded as part of short-read sequencing analyses (Audano et al., 2019; Chaisson et al., 2015, 2019) and by providing breakpoint resolution of variants previously inferred by indirect read-pair or read-depth approaches (R. L. Collins et al., 2020). Second, LRS has enabled the routine generation of genome assemblies (Koren et al., 2017; Shafin et al., 2020), and recent advances in sequencing technology and methods are now routinely producing phased genome assemblies fully capturing both haplotypes (Cheng et al., 2021; Lorig-Roach et al., 2023; Porubsky et al., 2021). These advances have begun to improve our understanding of mutational processes, recurrent mutations, and new variants associated with disease and adaptation (Begum et al., 2021; Dutta et al., 2019; Hsieh et al., 2021; Miller et al., 2022; Porubsky et al., 2022).

Consequently, large-scale LRS efforts have enabled the construction of improved reference genomes including pangenomic representations of species (Liao et al., 2022) and exploration of the pattern of normal and disease variation across a variety of NIH initiatives in unprecedented detail, e.g., the *All of Us* (All of Us Research Program Investigators et al., 2019) and GREGoR (Chadwick & Chris Wellington, n.d.) programs. A critical question in such large-scale projects is the tradeoff between sensitivity and specificity for variant discovery as a function of genome coverage. This is especially important given that throughput and cost are still major limitations of LRS. In this study, we attempt to address this issue by comparing two of the most common platforms, Oxford Nanopore Technologies (ONT) and PacBio HiFi sequencing, as well as commonly used read-based and assembly-based variant callers. To establish a truth set for comparison, we analyze two deeply sequenced human genomes, HG00733 and HG002, with a specific focus on the recovery of SVs. Realizing that both LRS technologies and variant callers are under continuous development, this analysis is a snapshot in time that aims at informing experimental design to achieve high sensitivity and specificity within realistic economic boundaries.

## RESULTS

Because LRS data can enable phased *de novo* assembly, we distinguish two LRS approaches for variant discovery: read-based and assembly-based methods. We define read-based methodologies as those requiring alignment of individual sequencing reads to a reference genome and applying specific read-based variant-calling algorithms to these alignments to identify variants. Assembly-based methods, in contrast, first generate *ab initio* a whole-genome assembly from LRS reads without guidance from a particular reference genome, and then proceed analogously by aligning this assembly to a reference genome to call variants using assembly-based calling algorithms. Many different tools implement variant-calling algorithms and they differ in their support for sequencing technologies (PacBio, ONT, etc.), variant types (SVs, indels, etc.), or data input (assembly, reads, etc.). In this study, we limit our analysis to eight read-based callers: Clair3 [v0.1-r11] (Zheng et al., 2021), cuteSV [v1.0.13] (Jiang et al., 2020), DeepVariant [v1.3.0] (Poplin et al., 2018), Delly [v1.0.3] (Rausch et al., 2012), PEPPER-Margin-DeepVariant [r0.8] (Shafin et al., 2021), Sniffles [v2.0.2] (Smolka et al., 2022), PBSV [v2.8.0] (*Pbsv: Pbsv - PacBio Structural Variant (SV) Calling and Analysis Tools*, n.d.), and SVIM [v1.4.2] (Heller & Vingron, 2019), and two assembly-based callers: PAV [v1.2.2] (Ebert et al., 2021) and SVIM-asm [v1.0.2] (Heller & Vingron, 2020). Assemblies were generated considering three algorithms: hifiasm [v0.16.1] (Cheng et al., 2021), PGAS [v14-dev] (Ebert et al., 2021; Porubsky et al., 2021), and Flye [v2.9] (Kolmogorov et al., 2019).

We set out to determine how variant-calling performance differs depending on the platform, depth of sequence coverage (X), and computational method. For this assessment, we generated downsampled sets of HiFi and both standard and ultra-long ONT (UL-ONT) sequence data at depths of 5, 8, 10, 12, 15, 17, 20, 25, and 30X assuming a 3.1 Gbp haploid genome size. We applied standard practice algorithms and procedures and evaluated precision and recall of each algorithm for single-nucleotide variants (SNVs), small (<50 bp) indels (insertions and deletions), and SVs with respect to the human reference genome GRCh38. We consider two publicly available human genomes that have been sequenced extensively: HG002 (the Genome in a Bottle [GIAB] Ashkenazim child reference genome) (Wagner et al., 2022) and HG00733 (a Puerto Rican reference genome from the 1000 Genomes Project). In addition to GIAB analysis of HG002 (Zook et al., 2016), both genomes have been extensively characterized for genetic variants by both the Human Genome Structural Variation Consortium (HGSVC) (Ebert et al., 2021) and Human Pangenome Reference Consortium (HPRC) (Liao et al., 2022), which has led to the availability of thoroughly vetted variant callsets (Ebert et al., 2021) that are used in this study as truth sets (referred to as HGSVC Freeze 4). Both genomes have the advantage that they are targets of telomere-to-telomere (T2T) assembly development (Rautiainen et al., 2022) and, as such, more accurate and complete variant callsets will likely be available in the future to further refine truth sets for comparison. As both of these genomes have been characterized in multiple LRS efforts, sufficiently deep and high-quality input sets are available from both ONT and PacBio. For PacBio HiFi, these sets include 78.6X/17.9 kbp (depth/N50) and 99.54X/20.6 kbp for HG002 and HG00733, respectively. ONT standard length datasets were 153.4X/30.23 kbp and 92.3X/33.6 kbp and the UL-ONT data were 33.15X/96.4 kbp and 38.11X/132.7 kbp for HG002 and HG00733, respectively (**Supplemental Table S1**).

### Read-based variant calling

Read-based SNVs were called with DeepVariant and Clair3 and showed the least variability between callers and technologies out of all three variant categories. At sequence read depth below 15X, recall of PacBio HiFi-tuned algorithms consistently outperformed ONT by an average of 0.03 (**Figure 1**). In fact, at ∼10X coverage (current production from a single Sequel II SMRT cell) both precision and recall for HiFi data plateau while reaching a precision of 0.96 and recall of 0.90. At 5X coverage, DeepVariant and Clair3 showed on average 0.05 higher F1 scores in PacBio compared to ONT (**Supplemental Table S2**). This was demonstrated in both precision and recall with DeepVariant performing better with respect to precision and Clair3 with respect to recall. At coverage depths above 15X, the F1 score plateaued around 0.94 with recall being consistently higher than precision for all callers and technologies. The data suggest that HiFi is generally better with regard to recall but that 12X standard ONT and HiFi perform comparably. It should be noted that SNV calling for HG002 performed by GIAB has been subjected to extensive QC and specific regions are likely undercalled as reflected in the clustering of SNVs only observed in the LRS callsets. These clusters correspond to large blocks of highly identical segmental duplications, tandem repeats, and subtelomeric repeats (**Supplemental Figure S1**). In our analysis of 30X coverage datasets, we observe 639,007 SNV calls, which were not seen in GIAB for HG002. Of these 639,007 calls, 284,760 (44.56%) were observed by both ONT and HiFi suggesting true positive calls, though missing from the current GIAB set. This may help explain the precision plateaus at 90% across technologies and algorithms.

**Figure 1.**
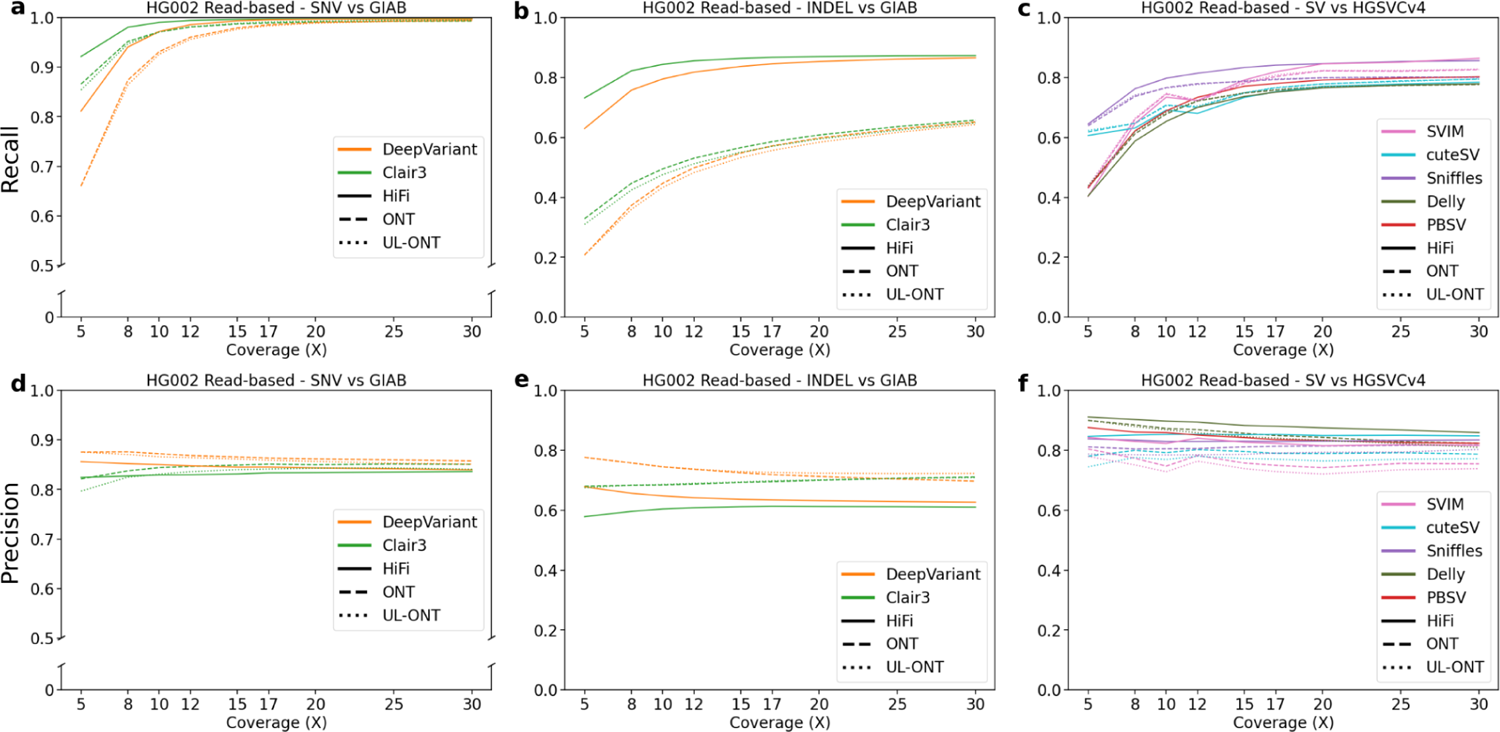
Precision and recall for variant classes as a function of LRS coverage using read-based algorithms for HG002. **a)** Recall of genome sample HG002 against Genome in a Bottle (GIAB) truth sets plotted against sequencing coverage for read-based callers Clair3 and DeepVariant. Clair3 with PacBio HiFi reaches the earliest recall plateau, while all callers show saturation by 20X. **b)** Recall against GIAB truth sets plotted against sequencing coverage for read-based callers across all algorithms capable of calling indels. Recall of both Clair3 and DeepVariant HiFi sets outperform their ONT counterparts. **c)** Recall against HGSVC truth sets plotted against sequencing coverage for read-based callers across all algorithms capable of calling structural variants (SVs). **d)** Precision as a function of sequence coverage. Single-nucleotide variant (SNV) precision remains flat beyond 10X, demonstrating the ability of callers to distinguish sequencing error from true SNVs. **e)** Precision plotted against sequencing coverage for read-based callers across all algorithms capable of calling indels. Precision values for all technologies and coverages remain flat, but here the increased precision of ONT callers is demonstrated. **f)** Precision plotted against sequencing coverage for read-based callers across all algorithms capable of calling SVs.

Indels, defined here as insertions or deletions less than 50 bp, show a similar profile. There is, once again, a characteristic plateau in F1 score around 12X sequence coverage. The greatest difference in recall is demonstrated in this subset between the HiFi and ONT platform (based on the R9 nanopore technology) (**Figure 1**). While precision remains high for ONT parameterizations of DeepVariant and Clair3 with an average of 0.82 across all measured depths, recall is noticeably lower when compared to PacBio HiFi reads, on average 0.39 less at depths less than or equal to 12X and 0.31 above 12X (**Supplemental Table S3**). Interestingly, for this class of variant, ONT reads prepared with standard library prep consistently outperform their UL counterparts with respect to precision. We observe a mean precision difference of 0.03 at or below 12X and a 0.07 difference above 12X in favor of standard ONT. Overall, recall for indels is higher in HiFi datasets at all coverages, while ONT callers are more precise. A large amount of community development has gone into refining variant callers for ONT and has allowed these callsets to reduce noise inherent to less accurate ONT sequence reads.

For SVs, we consider only insertions and deletions greater than or equal to 50 bp. SVs show the least variability between technologies (F1 standard deviation of 0.01 between HiFi and ONT sequencing platforms (**Supplemental Table S4**)). Both sequencing platforms and various coverages converge on a set of ∼12,800 SVs with each calling on average 25,634 SVs (**Figure 2**). Different read-based callers, however, show considerable variation. While recall remains low at lower sequence depth, mainly due to random sampling bias, two callers stand out as having the greatest precision: PBSV and Delly. Both callers consistently perform with high precision (mean 0.89) at low coverage depths and remain consistently high as depth increases. However, this does come with the above-mentioned tradeoff between precision and recall. As one increases, the other will decrease. In terms of recall at low-coverage sequence read depths below 12X, Sniffles performs best with a mean 0.63/0.84/0.71 precision/recall/F1 with cuteSV a close second (0.57/0.84/0.67).

**Figure 2.**
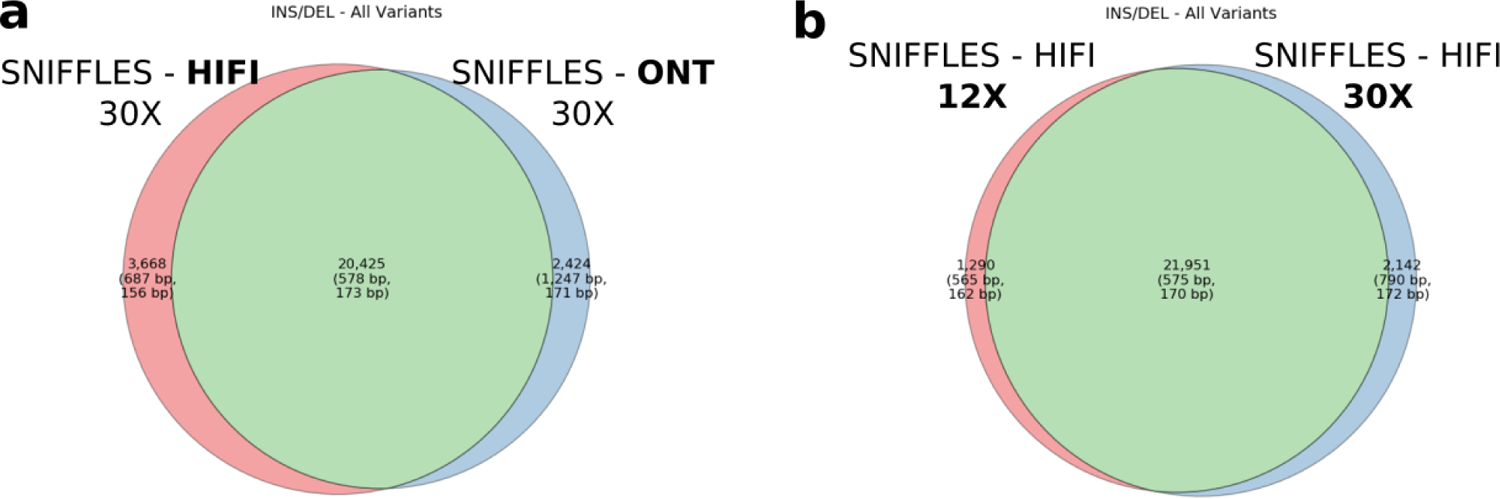
SV discovery. **a)** Venn diagram comparing Sniffles detection of SVs (both insertions and deletions) for 30X HiFi and 30X standard ONT input readsets. Of the variants unique to one technology or the other, 85% map to tandem repeat regions, which suggests breakpoint resolution rather than technology-specific bias is driving the difference. **b)** Venn diagram comparing Sniffles SV discovery at 12X and 30X HiFi callsets. A consistent set of calls is generated above 12X.

### Assembly-based variant calling

Assembly-based callers have the advantage that they call variants from large contiguous haplotype blocks essentially providing access to larger and more complex forms of genetic variation and providing extended phasing for all forms of genetic variation (Wagner et al., 2021). We generated assemblies using three algorithms: hifiasm (v0.16.1), PGAS (v14-dev), and Flye (v2.9) where applicable. Hifiasm and PGAS assemblies were generated for the PacBio HiFi readsets, and Flye assemblies for the ONT reads. All variants were called using the phased assembly variant (PAV) caller (Ebert et al., 2021) in addition to SVIM-asm specifically for SVs. It should be noted that the state of genome assembly for HiFi and ONT are not easily comparable. While HiFi reads can be assembled with numerous algorithms and assessed for phasing accuracy, ONT reads provide a greater challenge due to higher sequence error and fewer algorithms that combine both assembly and phasing. Methods such as Shasta (Shafin et al., 2020), wtdbg2 (Ruan & Li, 2020), and Canu (Koren et al., 2017) show considerable promise, but currently contiguous, haplotype-phased assemblies are not as easily generated and thus have not been utilized as frequently in current studies.

SNV calling with assembly-based callers generally underperforms read-based discovery especially at lower coverages. Precision in ONT and UL-ONT assembly-based methods shows the greatest difference with an average reduction of 0.33 across all sequencing depths (**Figure 3**). This is especially true in low-coverage (<12X) scenarios, and is driven by an excess of assembly-based SNV calls in ONT datasets (mean 8.33M in ONT; mean 10.00M in UL-ONT). PacBio HiFi methods have the opposite problem in that they underreport SNVs with a mean of 3.00M calls, although that does not greatly affect precision. This undercalling in HiFi assembly-based SNV callsets is a result of far less of the genome being assembled into haplotype-resolved contigs at lower coverages (**Figure 4**). However, when coverage reaches 12X, assembly-based methods show excellent recall (mean 0.96) for SNVs across all technologies (**Supplemental Table S5**) which mirrors the plateau observed in read-based methods. Below this threshold, read-based callers recall nearly 4X more (2,551 vs. 651) SNV windows based on recovery of over 90% of variants partitioned into 1 Mbp (**Figure 4**). Overall, SNV calling in low (less than 12X) coverage assemblies is not recommended, but coverages at or above 12X provide comparable precision and recall as their read-based counterparts with an average of 0.02 lower recall and 0.10 lower precision.

**Figure 3.**
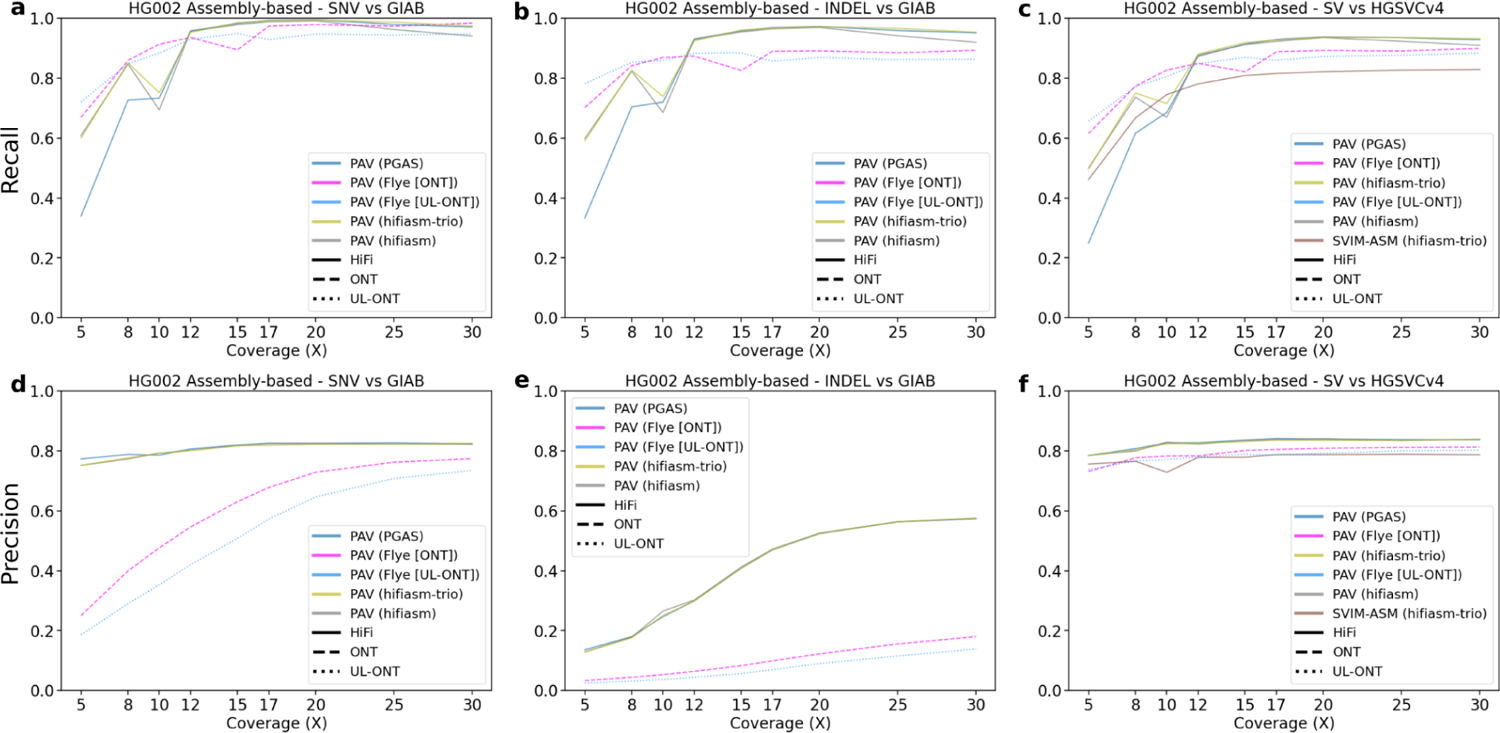
Precision and recall for variant classes as a function of LRS coverage using assembly-based algorithms for HG002. **a)** Recall for HG002 for GIAB truth sets plotted against sequencing coverage for assembly-based callers across all algorithms capable of calling SNVs. **b)** Recall for HG002 against HGSVC truth sets plotted against sequencing coverage for assembly-based callers across all algorithms capable of calling indels. Recall in ONT assemblies performs better at low coverages before being surpassed by HiFi assemblies at 12X. **c)** Recall for HG002 against the HGSVC Freeze 4 truth set plotted against sequencing coverage for assembly-based callers across all algorithms capable of calling SVs. **d)** Precision for HG002 against HGSVC truth sets plotted against sequencing coverage for read-based callers across all algorithms capable of calling SNVs. ONT methods are comparable to HiFi precision at high coverages but are noticeably worse at coverages below 15X. **e)** Precision plotted against sequencing coverage for assembly-based callers across all algorithms capable of calling indels. Like read-based methods, values for all technologies and coverages remains low, likely due to the incomplete nature of indels in complex regions in the GIAB truth set. **f)** Precision plotted against sequencing coverage for assembly-based callers across all algorithms capable of calling SVs.

**Figure 4.**
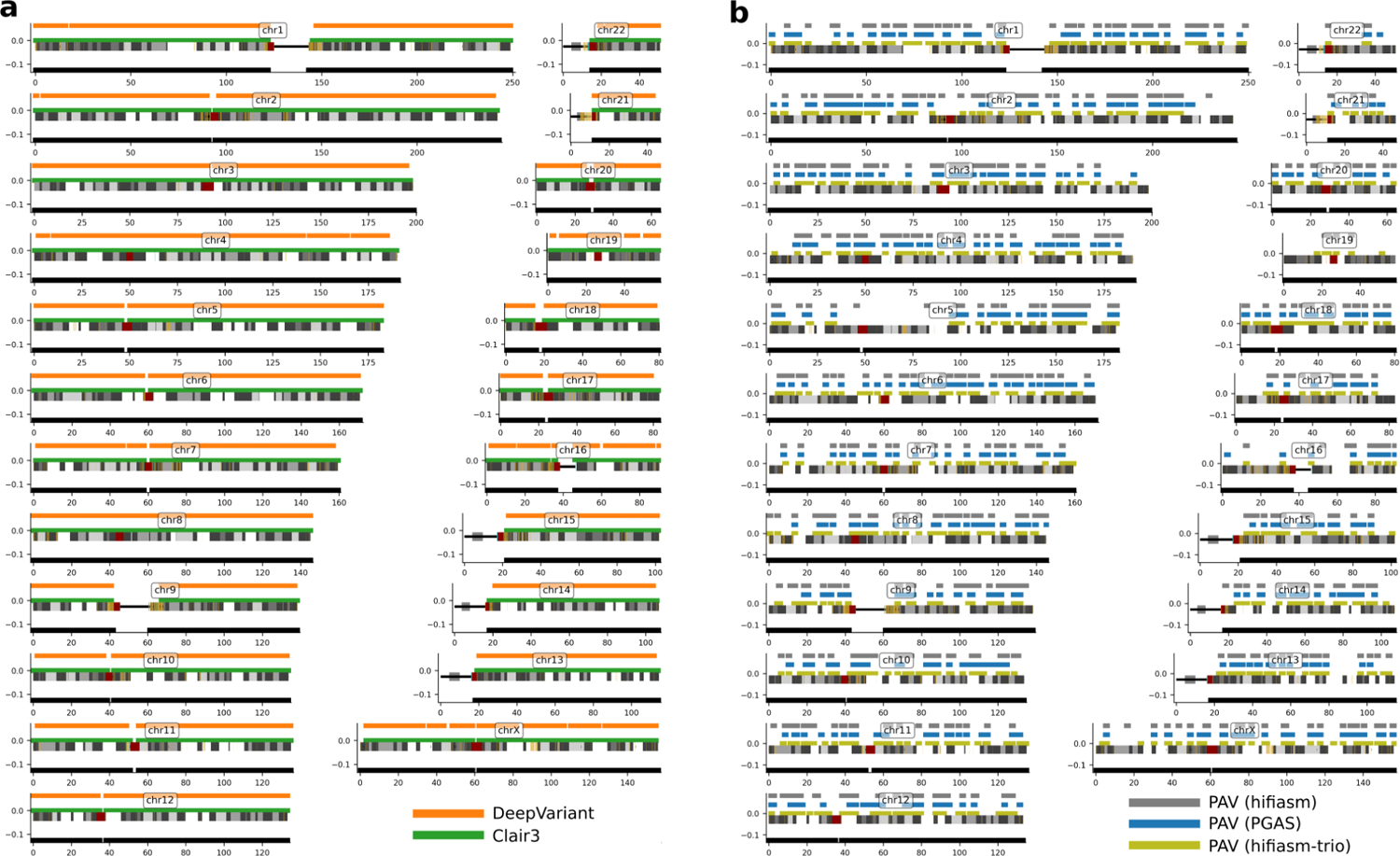
Ideogram comparison of autosomal SNV recall at 8X for PacBio HiFi. **a)** PacBio HiFi (8X) read-based recall of HG002 SNVs against GIAB truth sets. A bar over a chromosome depicts a 1 Mbp window where there was >90% SNV recall for Clair3 (green) and DeepVariant (orange) with all regions where SNV were called in black. **b)** PacBio HiFi (8X) assembly-based recall of HG002 GIAB SNV truth set using PAV. There are fewer 1 Mbp windows with >90% recall irrespective of assembly algorithm including hifiasm-trio (yellow), PGAS (blue), or hifiasm (gray). The black bar under the chromosome represents 1 Mbp windows with SNVs in the truth set.

Detecting indels from assembly-based methods is especially challenging (**Figure 3**), in part due to the known LRS error profiles associated with indels of smaller motif sizes (Delahaye & Nicolas, 2021; Wenger et al., 2019). Inability to correct these errors at low sequencing depth significantly inflates indel counts (1,145,880 indel insertion calls on average in PacBio HiFi 5X vs. 444,045 indel insertion calls in PacBio 30X). As such, precision is lowest for indels called in assemblies below 12X (**Supplemental Table S6**). In ONT datasets, this issue is exacerbated by an order of magnitude at reduced coverages (8,105,758 at 5X) and remains problematic even at high coverage (1,137,763 at 30X). Precision estimates, however, may be underestimated due to the limited capability of Illumina to detect variation in more complex regions of the genome that were not accessible to the GIAB truth set. Additional development and orthogonal validation of indels should be an active area of LRS technology development.

SVs follow the trend of assembly-based callsets in general with a steep recall curve, steady precision curve, and early plateau across sequencing depths and technologies. For low (below 8X) HiFi coverages, assembly-based methods underperform their read-based counterparts with respect to recall by an average of 0.03 (**Supplemental Table S7**). While ONT assemblies demonstrate higher recall than their read-based counterparts by 0.09 and 0.10 for standard ONT and UL-ONT, respectively. Above this coverage, all assembly-based methods outperform read-based methods by at least 0.08 for recall. The HG002 assemblies using PacBio HiFi reads at 10X sequencing depth are a clear outlier and may be attributable to a systematic failure to remove false duplications. We did not observe a similar outlier in HG00733. Although the assembly size is larger than expected, metrics such as contiguity (N50) and callable loci are consistent with other assemblies. Similar outliers may be avoidable with deeper coverage to support high-quality assembly-based callsets (Ebert et al., 2021; Liao et al., 2022).

### Cross-callset comparisons

Because LRS technologies claim to access more of the genome and more complex classes of genetic variants, we first evaluate genome-wide SV callability. To assess callability across the genome, we first divided GRCh38 into 1 Mbp windows and intersected those windows with the HGSVC SV truth set for HG00733, yielding 2,679 and 2,482 windows for insertions and deletions, respectively. In order for a window to be established as callable, >90% of the calls contained in this window must be accurately recovered (**Figure 5**). At low coverages (5X), read-based methods outperform assembly-based methods for each respective technology. At these low coverages, Sniffles used with HiFi reads performs the best, recovering 1,118/2,482 (45%) windows when considering deletion calls. This is almost double the PacBio HiFi callable windows for assembly-based methods. This trend holds for insertions, but we do note that Flye assembly-based methods using UL-ONT perform better than Sniffles on HiFi reads. At 10X and above, the pattern switches with HiFi assembly methods outperforming all read-based callers with the starkest difference occurring at 15X where assembly-based methods recover an additional 500 Mbp and 383 Mbp of the genome (for insertions and deletions, respectively) than read-based methods.

**Figure 5.**
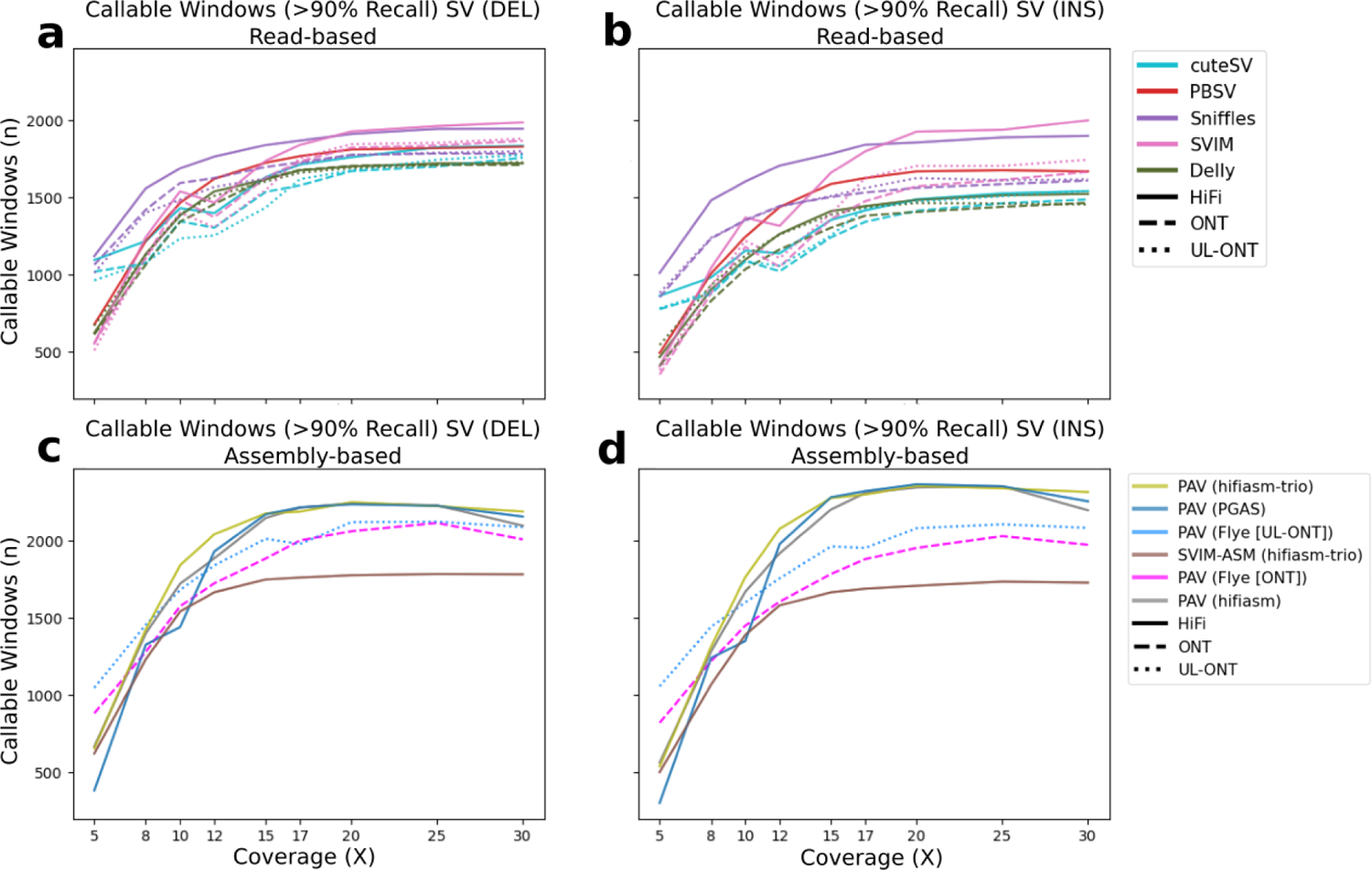
Evaluation of SV callable bases by technology and algorithm. Read-based callable windows for **(a)** insertions and **(b)** deletions, and assembly-based callable regions for **(c)** insertions and **(d)** deletions. Regions were compared against the HGSVC HG00733 truth set in 1 Mbp windows requiring at least 90% recall.

### Clinical SVs in HG002

A list of clinically relevant SVs was released for the GIAB sample HG002 (Wagner et al., 2021) including 273 challenging genes or regions that map to repetitive and structurally complex polymorphic regions. At 30X coverage, PBSV was able to recover 97% of these SVs in clinically relevant genes (**Supplemental Table S8**). However, at the lowest coverage depths, Sniffles, once again, drastically outperformed the other callers across all technology types, but especially with PacBio HiFi reads where it reports recall of 0.87 and 0.82 for SV insertions and deletions, respectively, at just 8X sequencing coverage. Compared to read-based methods, assembly-based methods demonstrated lower recall at low coverages with a max of 0.72 for insertions and 0.79 for deletions using Flye with UL-ONT and hifiasm (non-trio binned), respectively (**Supplemental Table S9**).

### Tandem repeat characterization

LRS technologies allow for more robust characterization of tandem repeats (Chaisson et al. 2015; Chaisson et al. 2019; Pendleton et al. 2015; Sedlazeck et al. 2018), the largest of which are known as variable number of tandem repeats (or VNTRs). After SNVs, tandem repeat variants are among the most abundant forms of human genetic variation comprising >20% of indels and >50% of SVs (Ebert et al., 2021) (**Supplemental Table S10**). Excluding these regions from analysis has little effect on recall, indicating that even though these regions have been difficult to characterize in prior studies, most LRS technologies and algorithms are able to detect these variants despite ambiguity in defining the exact breakpoints. However, inclusion of these regions potentially comes with a tradeoff in precision, particularly with read-based methods where we saw precision increase when we exclude tandem repeats at a consistent rate of 0.04 compared to precision in assembly-based callers, which were more precise in lower coverage scenarios with tandem repeats included (**Supplemental Figure S2**). This indicates that even at low coverages, assembly-driven variant calling can characterize such variation.

### Performance in homopolymer DNA

Accurately calling variants in homopolymer runs is challenging for both PacBio HiFi and ONT technologies (G. A. Logsdon et al., 2020; Mc Cartney et al., 2021; Shafin et al., 2021). These nonrandom error profiles impact precision and recall, especially for indel variant calls. When comparing the difference between all indel calls annotated with and without homopolymers, ONT callsets display a large difference between homopolymer and non-homopolymer DNA sequence precision and recall (**Supplemental Figure S3**). Even at high coverages, recall for insertions in homopolymer sequence is as much as 0.10 lower than when compared against the whole set. Notably, the effect that these sequence types have on precision even at higher depths is still prevalent with even 30X read-based methods showing a decrease of 0.06 between these regions. DeepVariant calls for UL-ONT reads show a decrease in homopolymer precision as sequencing depth increases. This could be due to a prior lack of training data with a ground truth for complex genomic regions uniquely aligned by this technology.

### Large variant discovery

Large (>10 kbp) SVs, especially insertions within or near repeat regions, frequently evade Illumina detection (Medvedev et al., 2009). An advantage of LRS technologies is that these events can be detected directly from the sequence of the reads or the assembly themselves. We assessed each method’s ability to recover large variants using the HGSVC validation set from HG00733 including 63 deletions and 40 insertions. For HiFi reads, two trends emerge: their limitation in detecting large insertions compared to ONT reads and their increased recall when assembled even at low coverages. HiFi reads consistently lag behind their ONT counterparts for large insertions, recovering only half of the insertions in standard ONT callsets and a third of the insertions detected in UL-ONT (**Supplemental Table S11**). However, by assembling these reads, HiFi datasets outperform ONT when sequence coverage exceeds 8X. Among read-based methods, UL-ONT performs the best with a minimum of 21/63 large deletions and 15/40 large insertions detected even at low sequence coverages (5X). Across all read-based algorithms, Sniffles recovers the greatest number of large events with a maximum of 0.67 and mean of 0.51 recall over all input types and coverages followed by cuteSV with 0.65 and 0.41, respectively. It should be noted that Delly failed to call any SVs above 10 kbp. HiFi assembly-driven methods perform the best overall with a maximum large variant recall of 0.87 and a mean of 0.65 when PAV is used (**Supplemental Table S12)**. Finally, it should be noted that both read-based and assembly-based methods recovered the largest (238 kbp) deletion, but only assembly-based methods identify the largest insertion of 51 kbp compared to the maximum event size in read-based methods of 32 kbp.

### ONT duplex reads and Revio HiFi data

PacBio and ONT are rapidly developing new sequencing technologies that improve LRS accuracy and throughput. For example, ONT recently released an improved flowcell (R10) as well as a new “duplex” sequencing method (*Oxford Nanopore Tech Update: New Duplex Method for Q30 Nanopore Single Molecule Reads, PromethION 2, and More*, n.d.) significantly improving individual read accuracy by sequencing both forward and complementary strands from the same single molecule (Sanderson et al., 2023). The new release of the Revio system from PacBio, in contrast, significantly increases throughput and affordability using a chemistry similar to that of the Sequel II platform (i.e., HiFi sequencing). The recent release of whole-genome sequencing (WGS) datasets from the GIAB sample HG002 allows these new emerging LRS platforms to be compared. We analyzed a 30X duplex dataset of WGS data released by ONT and compared precision and recall to standard ONT using R9.4.1 flowcells. We find that variant-calling recall for specific variant classes is substantially improved for duplex sequencing over R9 ONT variant calling at all sequence coverages and for all variant classes. The effect is most pronounced for indel recall at low coverage (≤10X) where duplex variant recall improves by 0.19 (**Figure 6**) when compared to standard ONT. Precision, however, is much more consistent with standard ONT methods. Of note, in our analysis, the precision of indel insertions actually diminishes when compared to standard ONT (an average of 0.06 reduction). This is possibly due to parameterization of variant-calling algorithms which have been largely adjusted for calling in a noisier, more error-prone, single-strand ONT signal.

**Figure 6.**
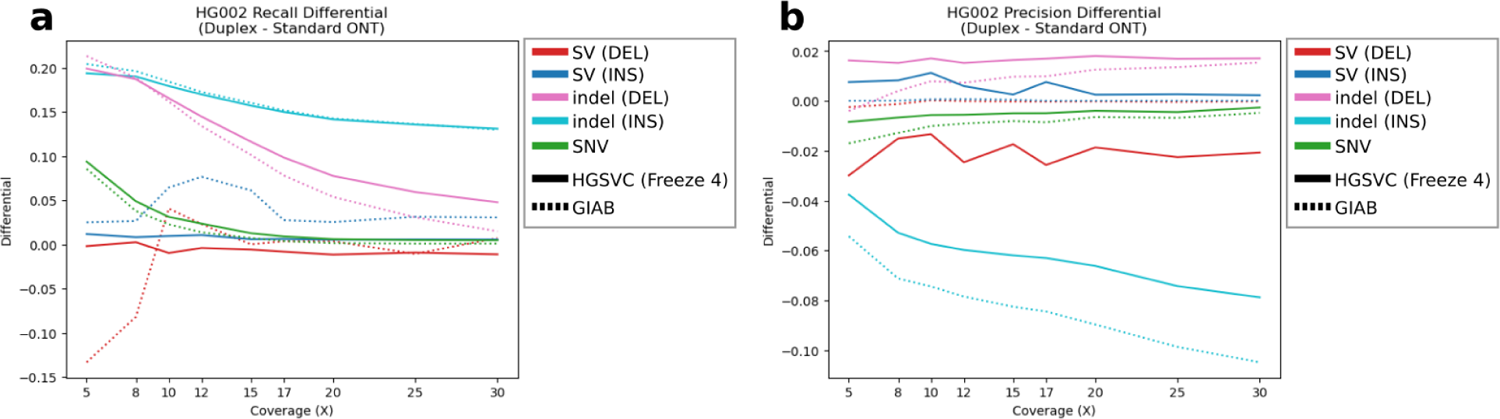
Comparison of precision and recall in duplex ONT variant calling versus standard ONT. Duplex ONT versus standard ONT variant calling **(a)** precision and **(b)** recall where anything above the y=0 line indicates an increase in performance compared to standard ONT and anything below the y=0 line indicates an decrease in performance compared to standard ONT.

Using 30X of WGS data from HG002 generated by the Revio system (*PacBio Revio*, 2022), we also constructed a phased human genome assembly using hifiasm. The results were nearly identical to an assembly produced from a Sequel II HiFi dataset, albeit with single flowcell. Both the contiguity (contig N50 = 44 Mbp [Revio] vs. 45 Mbp [Sequel II]) and accuracy (QV=57 [Revio] vs. 55 [Sequel II]) were virtually identical. Predictably, assembly-based variant calling were comparable for both recall (Pearson R = 0.984) and precision (Pearson R = 0.997) with some modest improvements in SNV recall (+0.02 vs. both truth sets) and small insertion precision (+0.06 vs. HGSVC Freeze 4) (**Table 1**).

**Table 1.** Revio versus Sequel II assembly-based callset comparison

| SAMPLE | SVTYPE | TRUTH SET | RECALL (HiFi) | RECALL (REVIO) | RECALL DIFF | PRECISION (HiFi) | PRECISION (REVIO) | PRECISION DIFF |
| --- | --- | --- | --- | --- | --- | --- | --- | --- |
| HG002 | SV (ins) | Freeze 4 | 0.94 | 0.91 | -0.03 | 0.823 | 0.865 | 0.042 |
| HG002 | SV (del) | Freeze 4 | 0.927 | 0.901 | -0.026 | 0.869 | 0.859 | -0.01 |
| HG002 | SNV | GIAB | 0.974 | 0.998 | 0.024 | 0.825 | 0.811 | -0.015 |
| HG002 | SNV | Freeze 4 | 0.97 | 0.992 | 0.022 | 0.897 | 0.879 | -0.018 |
| HG002 | indel (ins) | GIAB | 0.955 | 0.944 | -0.012 | 0.549 | 0.584 | 0.036 |
| HG002 | indel (ins) | Freeze 4 | 0.959 | 0.971 | 0.012 | 0.706 | 0.77 | 0.064 |
| HG002 | indel (del) | GIAB | 0.953 | 0.947 | -0.006 | 0.605 | 0.598 | -0.006 |
| HG002 | indel (del) | Freeze 4 | 0.965 | 0.986 | 0.022 | 0.776 | 0.789 | 0.014 |

\*PAV assembly-based variant-calling comparison for WGS data generated for HG002 on a Revio system compared to the 30X downsampled HG002 generated via the Sequel II platform compared to the HGSC truth set (Freeze 4) and Genome in a Bottle (GIAB).

## DISCUSSION

Within the limits of various algorithms and sequencing platforms analyzed here, we make a few general observations and recommendations based on our analysis against current truth sets (Ebert et al., 2021; Zook et al., 2016). With respect to SNV discovery, LRS coverage in excess of 12-fold begins to show a plateau with respect to sensitivity. Read-based approaches such as Clair3 (Zheng et al., 2021) and DeepVariant (Poplin et al., 2018) significantly outperform assembly-based detection methods, such as PAV, which have been geared to improve SV discovery and breakpoint definition (Audano et al., 2019; Ebert et al., 2021). While Clair3 with PacBio HiFi performs the best for SNVs, both deep convolutional network approaches (Clair3 and DeepVariant) show excellent recall with both ONT and PacBio above 20X sequence. Irrespective of the sequencing platform, sequence coverage at 8X or lower shows significant reduction in performance and is not advised for large-scale sequencing projects dedicated to variant discovery.

By contrast, all LRS platforms currently underperform for indel variant calling and, predictably, they perform the most poorly in regions of homopolymer runs as well as short tandem repeats— precisely the regions that are most mutable for this class of variation (Willems et al., 2014). Given that caveat, we would recommend PacBio HiFi read-based methods for recall (0.69 vs. 0.61) across all read coverages and ONT for precision, although the difference is slight (0.68 vs. 0.66 mean precision for ONT vs. HiFi, respectively). A major challenge facing human genetics is the existence of a well-vetted and complete truth set for indel variants—detailed studies over the years have restricted analyses to specific regions of the genome owing to the high rate of false positives and false negatives from more mutable and difficult-to-sequence regions (Krusche et al., 2019; Olson et al., 2022; Zook et al., 2019). Our results suggested that haplotype-resolved assemblies offer some improvement for recall (an average of +0.14 across all coverages). Completely sequenced and assembled genomes where T2T chromosomal assemblies are established along with vetted indel callsets by multiple sequencing technologies (e.g., Sanger, Illumina, ONT, and PacBio) will be required to develop a more comprehensive truth set of indels for benchmarking. Resources such as the Platinum pedigree (CEPH pedigree 14633) by Illumina will be particularly useful as they enable studying phased genome assemblies and variant calling in the context of transmission within families (Eberle et al., 2017).

Both ONT and HiFi PacBio excel at SV detection, routinely detecting >20,000 SVs and consistently calling the same variants when sequence coverage exceeds 12X (**Figure 3**). In fact, approximately 85% of SVs in 30X datasets that are unique to one platform over another map to tandem repeat regions but are in close proximity (<10 kbp) and their size overlap suggests that differences in alignment and breakpoint definition are still potentially more rate-limiting as opposed to platform differences in sensitivity. The advance of LRS for SV detection when compared to Illumina WGS has been well established over the years (Chaisson et al., 2015, 2019; Sedlazeck et al., 2018; Shafin et al., 2021) and more sophisticated callers as well as computational pipelines continue to be developed to discover and characterize SVs as part of routine callsets (Kolmogorov et al., 2023). While ONT, and especially UL-ONT, performed well for detecting large insertions (**Supplemental Table S11**), overall, assembly-based approaches (especially hifiasm) showed the greatest specificity and precision when calling large SVs (>50 kbp) (**Supplemental Table S12**). Because large SVs are much more likely to have phenotypic consequence and precise breakpoints are relevant to the effect of this consequence, assembly-based strategies should strongly be considered when applying LRS to solving cases of Mendelian and *de novo* disease (Miller et al., 2021). However, generation of phased genome assemblies requires deeper sequencing coverage (at least 15-20X) and, as such, is still a more expensive option.

In summary, when deciding LRS depth targets, the intended purpose of the project must be considered. If the goal is recovery and characterization of SNVs at a population scale, low-depth read-based methods will provide the right balance of maximizing discovery and number of samples in the study. However, if the goal is sequence resolution of large and complex variants at the level of individual patients, assembly-based methods, in particular hifiasm, are currently one of the most accurate strategies for building phased genome assemblies but require greater investment in terms of sequence coverage (well beyond 15X) and computational processing.

Importantly, the LRS platforms continue to rapidly evolve in terms of accuracy (ONT) and throughput (PacBio). Improved modeling of the platform-dependent errors as well as newer pores, or techniques (duplex sequencing) for ONT show considerable promise with suggestions that sequencing accuracy may in fact rival or surpass that of Illumina (Kolmogorov et al., 2023). Changes such as duplex sequencing with the R10 pore, however, currently come at a cost of lower throughput (Sanderson et al., 2023) and, as a result, added expense to achieve deep coverage. For the last three years, PacBio HiFi has dominated the field with respect to accuracy in large part due to the advent of circular consensus sequencing (CCS); however, multiple flowcells have been required to achieve deep sequence. The release of the new Revio platform earlier this year significantly increases throughput and decreases costs which will aid production of high quality and contiguous assemblies comparable to that of those generated previously by multiple Sequel II flowcells. Both platforms are currently highly complementary. Recently, algorithms that aim to incorporate the strengths of both PacBio HiFi and ONT reads to generate *de novo* T2T assemblies have shown very promising results (Rautiainen et al., 2022). Such hybrid technology approaches have the potential to supplant any single LRS technology as soon as the costs drop and the production of LRS assemblies become routine. The benefit of complete T2T variant discovery should not be underestimated.

## METHODS

### LRS datasets and data availability

#### ONT data generation

UL-ONT data were generated from the HG00733 lymphoblastoid cell line according to a previously published protocol (G. Logsdon, 2022). Briefly, 3-5 x 107 cells were lysed in a buffer containing 10 mM Tris-Cl (pH 8.0), 0.1 M EDTA (pH 8.0), 0.5% w/v SDS, and 20 ug/mL RNase A (Qiagen, 19101) for 1 hour at 37°C. 200 ug/mL Proteinase K (Qiagen, 19131) was added, and the solution was incubated at 50°C for 2 hours. DNA was purified via two rounds of 25:24:1 phenol-chloroform-isoamyl alcohol extraction followed by ethanol precipitation. Precipitated DNA was solubilized in 10 mM Tris (pH 8.0) containing 0.02% Triton X-100 at 4°C for two days. Libraries were constructed using the Ultra-Long DNA Sequencing Kit (ONT, SQK-ULK001) with modifications to the manufacturer’s protocol. Specifically, ∼40 ug of DNA was mixed with FRA enzyme and FDB buffer as described in the protocol and incubated for 5 minutes at RT, followed by a 5-minute heat-inactivation at 75°C. RAP enzyme was mixed with the DNA solution and incubated at RT for 1 hour before the clean-up step. Clean-up was performed using the Nanobind UL Library Prep Kit (Circulomics, NB-900-601-01) and eluted in 225 uL EB. 75 uL of library was loaded onto a primed FLO-PRO002 R9.4.1 flowcell for sequencing on the PromethION, with two nuclease washes and reloads after 24 and 48 hours of sequencing.

#### PacBio HiFi data generation

PacBio HiFi data were generated from the HG00733 lymphoblastoid cell line as previously described (G. A. Logsdon et al., 2021) with modifications. Briefly, DNA was extracted from 4.3×10^6 cells using the Monarch HMW DNA Extraction Kit for Cells and Blood (New England Biolabs) with 1400 rpm lysis speed. After UV absorption and fluorometric quantification (Qubit High Sensitivity DNA kit, Thermo Fisher) on the DS-11 FX instrument (Denovix) and evaluation of DNA integrity on FEMTO Pulse (Agilent), 12 μg of DNA was prepared for sequencing using Megaruptor 3 shearing (Diagenode, settings 19/31) and the Express Template Prep Kit v2 and SMRTbell Cleanup Kit v2 (PacBio). The library was size-selected on a PippinHT instrument (Sage Science) using a 15 kbp high-pass cut. Five SMRT Cell 8Ms were run on a Sequel II instrument using Sequel II chemistry C2.0/P2.2 with 30-hour movie times, 2-hour pre-extension, and adaptive loading targets of 0.8-0.85 (PacBio). Circular consensus calling was performed with CCS version 6.0.0 (SMRT Link v.10.1) and reads with estimated quality scores ≥Q20 were selected for downstream analysis.

#### Reference genome and reliable regions

To support long-read mapping, only the primary GRCh38 assembly was used, which includes chromosome scaffolds, the mitochondrial assembly, unplaced contigs, and unlocalized contigs. No alts, patches, or decoys were present in the assembly during the alignment stages. This reference was used previously (Audano et al., 2019; Ebert et al., 2021) and is available for download here: ftp://ftp.1000genomes.ebi.ac.uk/vol1/ftp/data_collections/HGSVC2/technical/reference/20200513_hg38_NoALT/. Whole-genome analysis was restricted to regions outside centromeres, pericentromeric repeats, and the mitochondrial chromosome where variant calls were previously determined to be less reproducible (Audano et al., 2019; Ebert et al., 2021). This is available here: http://ftp.1000genomes.ebi.ac.uk/vol1/ftp/data_collections/HGSVC2/technical/filter/20210127_LowConfidenceFilter/

### Downsampling

In-house python scripts were utilized to read in indexes for our input datasets and subsample reads randomly up to the desired threshold. We then used SAMtools fqidx to extract the desired reads from our larger sets and partitioned them into individual bins.

### Whole-genome alignment

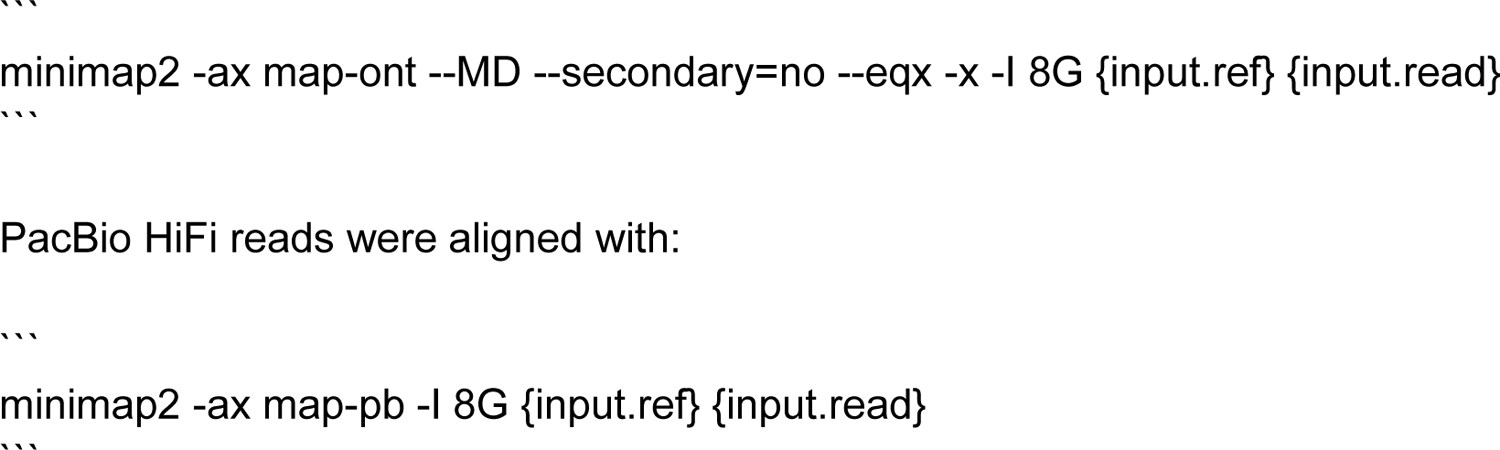

### Assemblies

We employed two approaches to generate phased whole-genome assemblies for all PacBio HiFi sampling depths: we used the PGAS pipeline as previously described (parameter settings v14-dev, (Ebert et al., 2021; Porubsky et al., 2021), hifiasm v0.16.1), which does not rely on parental data to derive genome-wide phase information. Additionally, we executed hifiasm v0.16.1 (Cheng et al., 2021) with default parameters in trio-binning mode, leveraging parental short reads to obtain phase information. For the ONT and UL-ONT readsets, we implemented a two-step process employing first the Flye assembler v2.9 (Kolmogorov et al., 2019) to generate unphased whole-genome assemblies with default parameters (preset “--nano-hq” and “--genome-size” of 3.1 Gbp). Next, these assemblies were converted into diploid assemblies using the HapDup v0.6 tool (Kolmogorov et al., 2019; Shafin et al., 2020) with default parameters (preset “ont”).

### Read-based variant calling

We used Clair3 [v0.1-r11] (Zheng et al., 2021) cuteSV [v1.0.13] (Jiang et al., 2020), DeepVariant [v1.3.0] (Poplin et al., 2018), Delly [v1.0.3] (Rausch et al., 2012), PBSV [v2.8.0] (*Pbsv: Pbsv - PacBio Structural Variant (SV) Calling and Analysis Tools*, n.d.), PEPPER-Margin-DeepVariant [r0.8] (Shafin et al., 2021), Sniffles2 [v2.0.2] (Smolka et al., 2022), and SVIM [v1.4.2] (Heller & Vingron, 2019) in order to call SVs from the aligned PacBio HiFi, ONT, and UL-ONT reads at the different coverage levels.

The commands used for each caller and technology are listed below:

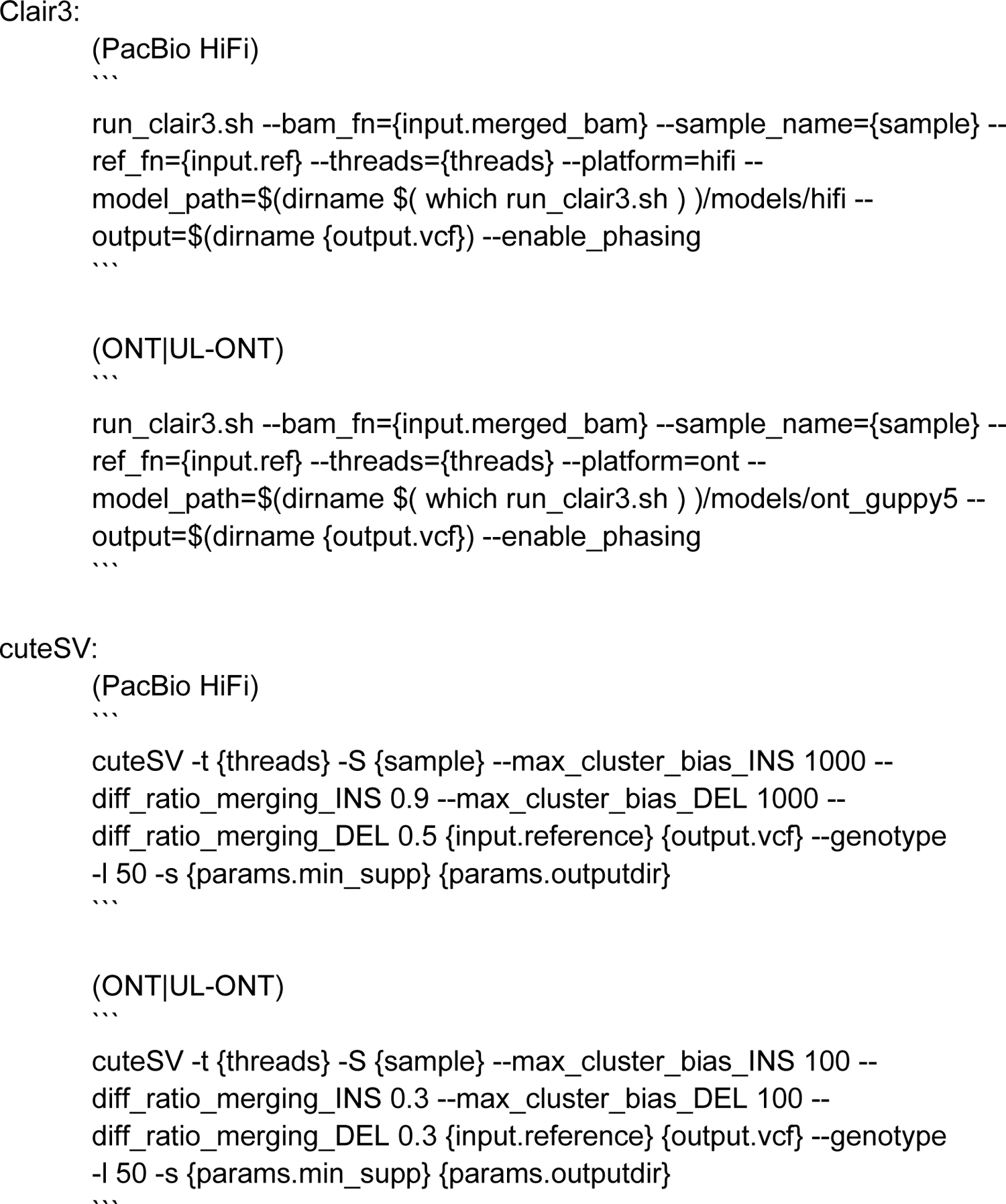

In addition, we filtered the cuteSV calls based on the minimum read support reported in the output VCF, as it generated unfiltered calls. Similarly, we filtered the SVIM calls based on the reported quality. In both cases, we used value 2 for coverages ≤5; 3 for coverages ≤10; 4 for coverages ≤20; 5 for coverages ≤25; and 10 for coverages >30. These values were selected such that they result in the highest F-scores when comparing the filtered calls to the GIAB medically relevant SVs for HG002. The pipeline used for SV calling with cuteSV, Sniffles2, and SVIM can be found here: https://github.com/eblerjana/lrs-sv-calling.

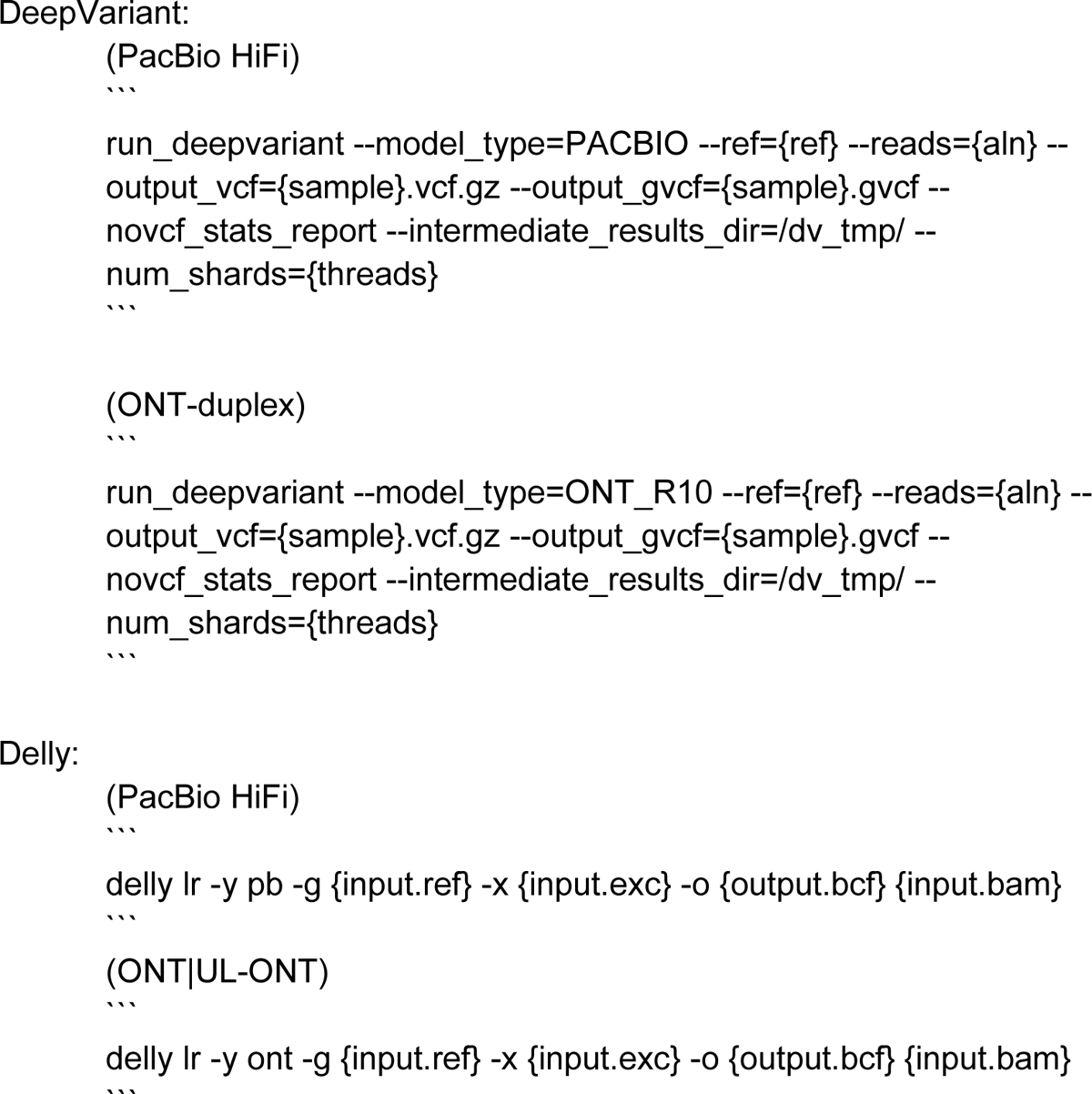

Excluded regions for Delly can be found here: https://github.com/dellytools/delly/blob/main/excludeTemplates/human.hg38.excl.tsv

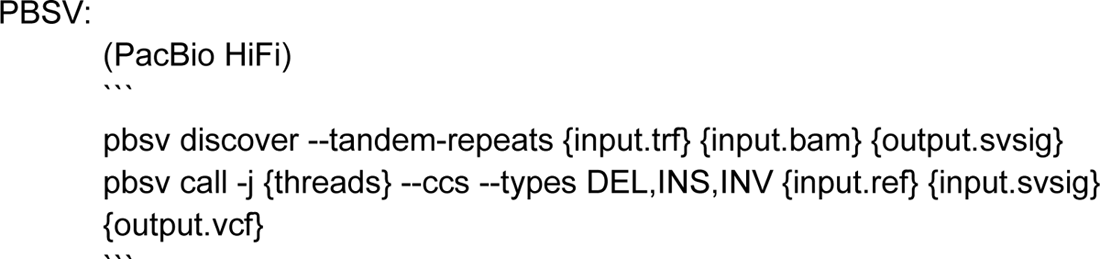
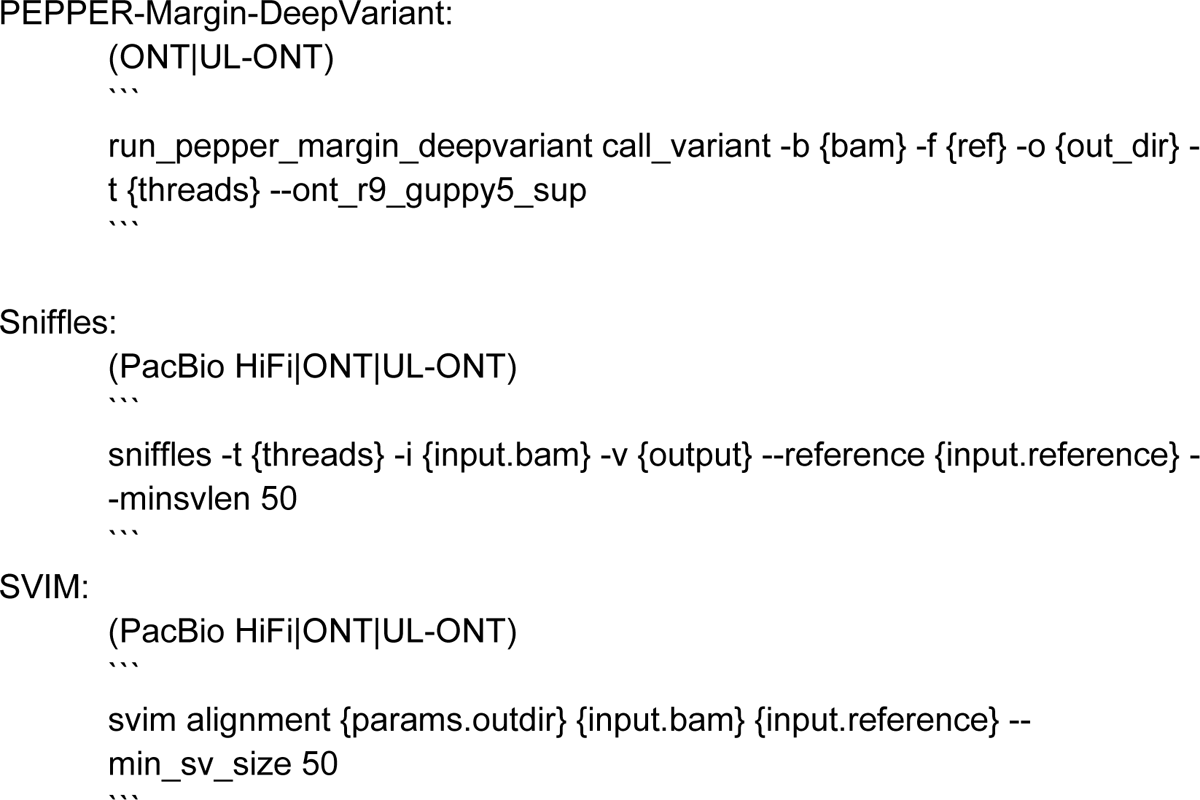

### Assembly-based variant calls

PAV (Ebert et al., 2021) was applied to phased assemblies using default parameters. Briefly, assemblies were mapped to the GRCh38 reference genome with minimap2 2.17 (Li, 2018), alignment trimming was performed to eliminate redundantly mapped bases, and variant calling was performed to detect variants within alignments as well as large SVs that fragmented alignment records into multiple parts.

### Variant merging and annotations

Variant call comparisons were performed using svpop. SNV-based comparisons were performed using the overlap feature nrid (nonredundant ID match), which requires variants to have the same SNV ID (#CHROM-POS-SNV-REF-ALT) to be called the same. Additionally, indels and SVs were matched using szro-50-200, which first matches variants on ID (#CHROM-POS-SVTYPE-SVLEN), then 50% reciprocal overlap, and then finally variants of the same type that are within 200 bp of each other and have reciprocal size overlap of 50%. This strategy allows for increased accuracy in complex regions of the genome where alignments can be biologically ambiguous. Reference-based annotations for genomic sequence content (e.g., homopolymer, TRF) are taken directly from the UCSC Genome Browser and the UCSC GoldenPath. This is a built-in functionality of SVPOP for GRCh38.

### F1 score

F1 score is defined as the harmonic mean between precision and recall and seeks to represent precision and recall in one metric.

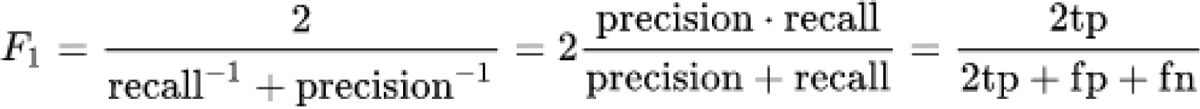

## DATA ACCESS

HG002 HiFi data was acquired as part of the HPRC and is available here: https://s3-us-west-2.amazonaws.com/human-pangenomics/index.html?prefix=T2T/scratch/HG002/sequencing/hifi/. HG002 ONT, UL-ONT, and duplex ONT data were acquired from the EPI2ME project (*EPI2ME^TM^*, n.d.). HG002 Revio data was acquired directly from PacBio and is available here: https://downloads.pacbcloud.com/public/revio/2022Q4/. HG00733 HiFi, ONT, and UL-ONT data were generated in house and have been submitted to the NCBI BioProject database (https://www.ncbi.nlm.nih.gov/bioproject/) under accession number PRJNA966152.

## ACKNOWLEDGMENTS

This work was supported, in part, by US National Institutes of Health (NIH) grants R01HG010169, U24HG007497, and U01HG010971 to E.E.E. Computational infrastructure and support were provided by the Centre for Information and Media Technology at Heinrich Heine University Düsseldorf. E.E.E. is an investigator of the Howard Hughes Medical Institute. This article is subject to HHMI’s Open Access to Publications policy. HHMI lab heads have previously granted a nonexclusive CC BY 4.0 license to the public and a sublicensable license to HHMI in their research articles. Pursuant to those licenses, the author-accepted manuscript of this article can be made freely available under a CC BY 4.0 license immediately upon publication.

## AUTHOR CONTRIBUTIONS

W.T.H. conducted assembly generation, variant calling, variant annotation, merging, and data analysis and visualization in addition to writing the text. P.E. produced assemblies and variant calls. J.E. produced variant calls. P.A.A. assisted with variant calling and variant merging. K.M.M. produced PacBio HiFi data for HG00733. K.H. produced ONT data for HG00733. D.P. helped with assembly analysis. C.R.B. provided structural guidance. T.M. provided assistance with evaluating precision and recall and experimental design. K.G. assisted with experimental design and caller parameterization. E.E.E. provided project oversight, biological insight, and major text additions. All authors read and approved the final manuscript.

